# ProtHGT: Heterogeneous Graph Transformers for Automated Protein Function Prediction Using Biological Knowledge Graphs and Language Models

**DOI:** 10.1101/2025.04.19.649272

**Authors:** Erva Ulusoy, Tunca Doğan

## Abstract

**Motivation:** The rapid accumulation of protein sequence data, coupled with the slow pace of experimental annotations, creates a critical need for computational methods to predict protein functions. Existing models often rely on limited data types, such as sequence-based features or protein-protein interactions (PPIs), failing to capture the complex molecular relationships in biological systems. To address this, we developed ProtHGT, a heterogeneous graph transformer-based model that integrates diverse biological datasets into a unified framework using knowledge graphs for accurate and interpretable protein function prediction.

**Results:** ProtHGT achieves state-of-the-art performance on benchmark datasets, demonstrating its ability to outperform current graph-based and sequence-based approaches. By leveraging diverse biological entity types and highly representative protein language model embeddings at the input level, the model effectively learns complex biological relationships, enabling accurate predictions across all Gene Ontology (GO) sub-ontologies. Ablation analyses highlight the critical role of heterogeneous data integration in achieving robust predictions. Finally, our use-case study has indicated that it’s possible to interpret ProtHGT’s predictions via exploring the related parts of our input biological knowledge graph, offering plausible explanations to build or test new hypotheses.

**Availability and Implementation:** ProtHGT is available as a programmatic tool on Github and as a web service at Hugging Face.

**Contact:** To whom the correspondence should be addressed: Tunca Doğan

## 1 Introduction

Functional annotation of proteins is essential for unravelling cellular processes, understanding disease mechanisms, and identifying potential therapeutic targets. Advances in high-throughput sequencing technologies have generated a massive amount of protein sequence data, presenting both vast opportunities and significant challenges. While experimental methods are considered the gold standard for determining protein function, they are time-consuming, expensive, and incapable of keeping pace with the rapid growth of data, leaving the majority of proteins in databases without functional annotations (Shehu, Barbará and Molloy 2016).

To address this challenge, computational methods for protein function prediction have become essential. These approaches utilise existing biological knowledge and extract meaningful patterns from diverse data types, such as sequences, domains, and interaction networks, to infer protein functions. By automating the annotation process, they offer a scalable and cost-effective alternative to experimental techniques while also enabling the generation of new hypotheses that can guide validation efforts and enhance our understanding of complex biological systems. A widely-used resource for these methods is the Gene Ontology (GO), which provides a standardised and machine-readable framework for defining protein functions (The Gene Ontology Consortium *et al*. 2023). GO organises functional information into three main categories: molecular function (MF), describing a protein’s biochemical activity; cellular component (CC), identifying its location within the cell; and biological process (BP), defining the broader biological objectives it contributes to. This hierarchical framework is structured as a directed acyclic graph (DAG), where parent-child relationships reflect increasing specificity.

Protein function prediction methods can be broadly categorised into feature-based and relation-based approaches. Feature-based methods analyse individual protein attributes, such as sequences, domains, motifs, and structural properties, and utilise well-annotated proteins to make predictions (Shehu, Barbará and Molloy 2016). Among these, sequence-centric methods are the most widely adopted (Yan *et al*. 2023). By focusing on properties such as amino acid composition, conserved motifs, and sequence alignments, these methods infer functions of uncharacterized proteins via utilising the evolutionary relationships between them and well-annotated proteins. Tools like BLAST (Altschul *et al*. 1990) and PFAM (Bateman *et al*. 1999) have been central to these efforts, aligning sequences to identify homologs based on the assumption that sequence similarity correlates with functional similarity. However, this assumption is unreliable in certain contexts, especially in the case of distant relationships, leading to annotation errors (Shehu, Barbará and Molloy 2016). Machine learning-based methods have since emerged to address these limitations, enabling the identification of subtle, functionally relevant patterns that conventional alignment methods often overlook (Kulmanov and Hoehndorf 2020).

Beyond sequences, domains and other structural motifs are also employed to predict protein functions, as they serve as powerful indicators of biological roles. Domains, structural and functional units within proteins, are analysed to transfer functional annotations between proteins with similar domain compositions or arrangements (Doğan *et al*. 2016; You *et al*. 2018; Rojano *et al*. 2022; Ulusoy and Doğan 2024). Secondary (e.g., helices and sheets) and tertiary structures add another dimension of information (Roy, Yang and Zhang 2012; Ma *et al*. 2022), as proteins with similar 3D folds often perform similar functions even if their sequences differ (Koonin and Galperin 2003). Recognising the complementary strengths of sequence- and structure-based approaches, hybrid models have emerged as an effective solution, integrating multiple data types to improve prediction accuracy (Zhang, Freddolino and Zhang 2017; Zhang *et al*. 2019; Cai, Wang and Deng 2020).

On the other hand, feature-based methods face significant limitations. Most proteins operate within complex networks of interactions, and focusing solely on individual attributes, such as sequences or structures, fails to capture the broader context of these relationships. Additionally, feature-based methods heavily rely on annotated homologs and motifs, which limits their generalizability to less-studied proteins or diverse organisms. These constraints emphasize the need for integrative approaches like graph-based methods, which can model the relational dependencies that influence protein functions.

Graph-based methods consider the broader biological context in which proteins operate. Graphs (or networks) provide an intuitive way to represent complex biological systems, where nodes correspond to entities like proteins or genes, and edges denote their relationships. These structures can be analysed using traditional graph-theoretic algorithms or their novel machine learning-based counterparts, such as graph neural networks (GNNs), which are designed to learn from the relationships encoded within the graph (Zhou *et al*. 2020). GNNs work by iteratively aggregating information from a node’s neighbours, updating its representation to reflect both local and global influences. A widely used type of GNN is the graph convolutional network (GCN), which extends the concept of convolution from grid-based data (e.g., images) to the irregular structure of graph data (Kipf and Welling 2016).

Protein function prediction has benefited from using protein-protein interaction (PPI) networks, where nodes represent proteins and edges capture their interactions (You *et al*. 2021). In these methods, nodes are initialised with features such as domain composition or sequence embeddings, which are refined through graph convolutional layers. This iterative process allows the model to integrate information from neighbouring nodes and uncover both direct and indirect interaction patterns essential for functional inference. Another approach leverages the hierarchical structure of GO terms. In these models, GNNs propagate information through the GO hierarchy, assigning features derived from sequence or evolutionary data to query proteins (Zhao, Liu and Wang 2022). This ensures predictions remain consistent with the biological relationships encoded in the GO structure. Additionally, some methods focus on amino acid interaction graphs derived from protein 3D structures (Gligorijević *et al*. 2021; Lai and Xu 2022). In these graphs, nodes represent amino acids, and edges denote spatial interactions within the protein fold. Features like physicochemical properties or embeddings from protein language models are refined through graph convolutional layers, enabling the identification of functional regions, such as catalytic or binding sites, which are less apparent from sequence data alone.

While graph-based methods have had a transformative impact, they also have notable limitations. Many rely on single-type networks, such as PPI networks or GO hierarchies, which only capture a fraction of the complex relationships present in biological systems. Even heterogeneous graph approaches often focus on a narrow range of connections, such as protein-GO annotations, direct protein interactions, or GO term hierarchies. This limited scope overlooks the rich interplay of biological factors, including disease-phenotype associations, drug/metabolite interactions, domain compositions, and signalling/metabolic pathway involvement of proteins, generally reflected as performance and interpretability-related issues.

To address these challenges, more comprehensive biological/biomedical frameworks, such as biological knowledge graphs (KGs), are needed to harmonise and integrate diverse data types and their relationships. By incorporating a wide range of relational connections, such as PPIs, protein-drug interactions, pathway membership information, and disease-phenotype links, KGs offer a holistic view of biological systems (Himmelstein *et al*. 2017; Doǧan *et al*. 2021; Chandak, Huang and Zitnik 2023). This integration can provide a richer understanding of the underlying biology and enable more accurate protein function predictions, overcoming the limitations of traditional graph-based approaches.

In this study, we introduce ProtHGT, a novel heterogeneous graph learning model designed to operate on an extensive biological graph for Gene Ontology (GO)-based protein function prediction. As part of this work, we constructed a comprehensive KG, called CROssBAR KG (Doǧan *et al*. 2021), from 14 diverse data sources, encompassing 9 types of biological entities and 17 types of interactions between them. ProtHGT utilises a heterogeneous graph transformer-based machine learning architecture to provide a deeper understanding of protein functions by learning from this integrative KG. By utilising attention mechanisms, ProtHGT identifies and prioritises the most influential connections within the graph, effectively capturing the unique characteristics of each node and edge type and their specific contributions to determining protein functions.

ProtHGT outperforms existing methods on benchmark datasets, such as CAFA3 and DeepHGAT, and its ablation analyses highlight the critical role of heterogeneous data integration in enhancing predictive accuracy. The model’s ability to refine node features into biologically coherent representations further emphasizes the value of combining diverse datasets and advanced graph-based architectures. The biological validity of ProtHGT’s predictions was demonstrated through a use case study, which also showcased its interpretability, allowing researchers to trace predictions to their underlying connections. This interpretability, combined with its potential to provide new insights into functional genomics, makes ProtHGT a valuable tool for researchers in bioinformatics, with future directions including extensions to broader applications such as protein-disease association prediction, drug-target interaction modeling, and drug discovery, further solidifying its utility for addressing complex challenges in biology and biomedicine. ProtHGT is available both as a programmatic tool on https://github.com/HUBioDataLab/ProtHGT and as an intuitive web service on https://huggingface.co/spaces/HUBioDataLab/ProtHGT.

## 2 Material and Methods

### 2.1 Dataset

#### Input Knowledge Graph

To build a comprehensive knowledge graph (KG) that offers a holistic view of protein functions, we integrated diverse biological and biomedical datasets, including entity types such as proteins, pathways, diseases, phenotypes, and drugs/compounds, along with their interconnections. Each dataset was carefully preprocessed and standardised to ensure compatibility and seamless integration. The majority of the entities and relationships were retrieved using our previously developed CROssBAR system (Doǧan *et al*. 2021), a tool designed to integrate data from multiple databases and generate query-specific knowledge graphs (https://crossbar.kansil.org). Human proteins available in CROssBAR were queried, and the resulting knowledge graphs were merged into a single, unified KG. This initial KG served as the foundation for subsequent data enrichment.

To expand its scope, proteins from organisms extensively studied in the literature and featured in widely recognised protein function prediction benchmarks were incorporated into the dataset. Pairwise orthology relationships among these proteins were retrieved from the Orthologous MAtrix (OMA) database (Altenhoff *et al*. 2021), expanding the dataset to cover 29 organisms. To ensure high-quality annotations, only proteins curated in the UniProtKB/Swiss-Prot database (UniProt Consortium 2023) were included.

The dataset was further enriched by incorporating additional biological entities from various databases to provide functional context. Domains and their associations with proteins were sourced from the InterPro database (Hunter *et al*. 2009), and domain-functional term relationships were retrieved from the InterPro2GO dataset. Gene Ontology (GO) terms and their hierarchical relationships were obtained from the Gene Ontology resource (The Gene Ontology Consortium *et al*. 2023). GO term annotations of proteins were acquired from the Gene Ontology Annotation (GOA) database (Huntley *et al*. 2015), which provides high-quality functional annotations based on experimental results, computational predictions, and literature curation. Annotations with the evidence code “Inferred from Electronic Annotation (IEA)” were excluded to ensure reliability. Lastly, EC numbers and their protein associations were integrated from the Expasy ENZYME database (Bairoch 2000).

To prevent data leakage between training and test sets, proteins with high sequence similarity were clustered using the UniRef90 dataset (Suzek *et al*. 2015). From each cluster, only a single representative protein was included in the final dataset, ensuring that proteins with more than 90% sequence similarity were excluded from appearing in both sets. An overview of the node and edge types in the final heterogeneous knowledge graph is shown in Figure 1b, with dataset statistics provided in Table 1 and 2. It is also important to note that we utilised other train-test datasets (e.g., CAFA3 and DeepHGAT) obtained from the literature mainly to compare our performance results with others. Details regarding these analyses are provided in section 2.4.

**Figure 1.**
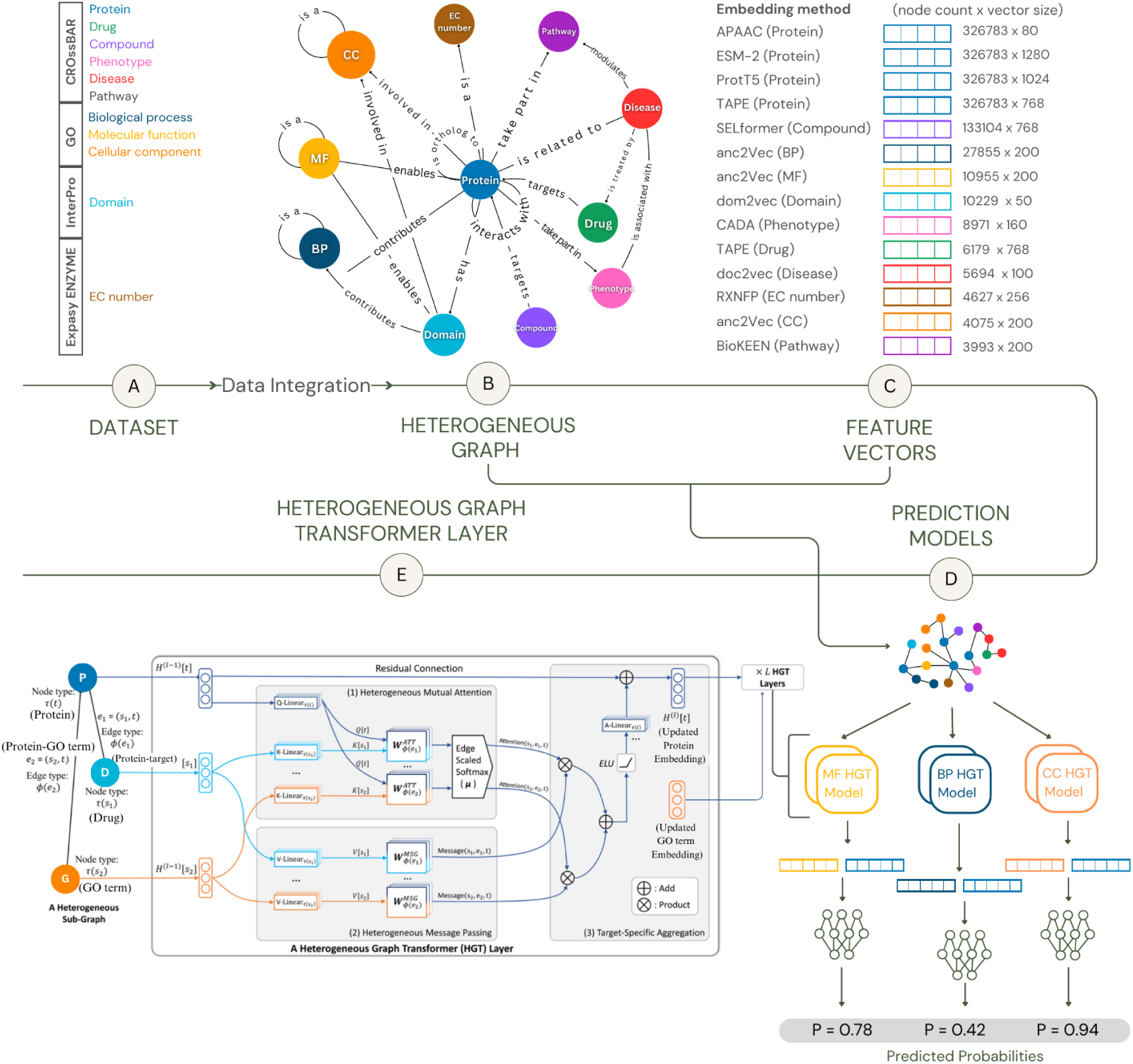
Overview of ProtHGT. **a)** The dataset integrates multiple biological data sources, including proteins, GO terms, and pathways. **b)** A heterogeneous graph is constructed to capture relationships between different node types. **c)** Feature vectors for nodes are derived using embedding methods specific to each data type. **d)** Prediction models are trained for Molecular Function (MF), Biological Process (BP), and Cellular Component (CC) using node embeddings. **e)** The Heterogeneous Graph Transformer (HGT) layer facilitates message passing and attention across node and edge types, enabling refined embeddings for improved prediction performance.

**Table 1.** Overview of node types and counts in training graph data.

| Node Type | Node Count |
| --- | --- |
| Protein | 326,783 |
| Compound | 133,104 |
| GO term | 42,885 |
| Domain | 10,229 |
| Phenotype | 8,971 |
| Drug | 6,179 |
| Disease | 5,694 |
| EC number | 4,627 |
| Pathway | 3,993 |
| **Total** | **542,465** |

**Table 2.** Overview of edge types and counts in training graph data.

| Edge Type | Edge Count |
| --- | --- |
| Protein-Biological process | 1,000,952 |
| Protein-Molecular function | 926,844 |
| Protein-Cellular component | 672,139 |
| Protein-Domain | 461,668 |
| Protein-EC number | 128,242 |
| Protein-Compound | 153,764 |
| Orthology | 125,372 |
| GO term-GO term | 83,831 |
| Protein-protein interaction | 82,524 |
| Protein-Pathway | 51,692 |
| Protein-Phenotype | 38,239 |
| Disease-Phenotype | 24,259 |
| Protein-Drug | 22,718 |
| Protein-Disease | 12,817 |
| Domain-GO term | 5,791 |
| Disease-Pathway | 1,682 |
| Disease-Drug | 298 |
| **Total** | **3,792,832** |

#### Node Features

To represent the inherent characteristics of each node type alongside their relational context, we employed state-of-the-art embedding methods known for their robust representational capabilities. These feature vectors served as the initial input to ProtHGT, enabling the model to capture both node-specific properties and complex relational patterns.

For protein nodes, we explored four types of embeddings, including traditional encoding approaches like amphiphilic Pseudo-Amino Acid Composition (APAAC) (Chou 2005) and pretrained protein language model embeddings such as ESM-2 (Lin *et al*. 2023), TAPE (Rao *et al*. 2019), and ProtT5 (Elnaggar *et al*. 2021). These methods were selected to explore the relative strengths of traditional approaches versus modern language model-based techniques in capturing the structural and functional nuances of proteins. While traditional encoding methods like APAAC offer a simpler, interpretable representation based on amino acid composition, language model embeddings are designed to capture deeper sequence-level dependencies and contextual information. GO term nodes were encoded using 200-dimensional embeddings produced by the anc2vec method (Edera, Milone and Stegmayer 2022). This approach captures three structural aspects of the Gene Ontology (GO) database: ontological uniqueness of terms, ancestral relationships, and sub-ontology classification. These embeddings were derived from the GO database version dated 2020-10-06.

Biotechnological (peptide) drug nodes were represented in a manner similar to protein nodes, using transformer-based embeddings derived from a pretrained protein language model (Rao *et al*. 2019). For domain nodes, 50-dimensional feature vectors were generated using the dom2vec method (Melidis and Nejdl 2021), which employs the word2vec algorithm (Mikolov *et al*. 2013) by treating protein sequences as sentences and domains as the words within those sentences. EC number nodes were represented using 256-dimensional embeddings derived from the SMILES representations (Weininger 1988) of their associated reactions, computed using the Python RXNFP library (Schwaller *et al*. 2021). Disease nodes were represented using 100-dimensional embeddings generated from their textual descriptions with the doc2vec method (Řeh uřek and Sojka 2010). Descriptions were primarily obtained from the PrimeKG dataset (Chandak, Huang and Zitnik 2023), and when unavailable, descriptions were retrieved from the original source databases where the nodes were obtained. In cases where no description was available, the disease names were utilised as input.

Biological pathway nodes were encoded as 200-dimensional embeddings generated using the TransE (Bordes *et al*. 2013) knowledge graph embedding method, applied to gene-biological pathway interaction networks via the Python BioKEEN library (Ali *et al*. 2019). For phenotypic term nodes, 160-dimensional embeddings were produced using the node2vec algorithm (Grover and Leskovec 2016) over gene-phenotype relationship networks, as implemented by the CADA tool (Peng *et al*. 2021). Lastly, compound nodes were encoded as 768-dimensional embeddings derived from their SELFIES representations using SELFormer (Yüksel *et al*. 2023), a pretrained chemical language model.

### 2.2 Architecture

The heterogeneous graph transformer (HGT) architecture is specifically designed to handle diverse types of nodes and edges in heterogeneous graphs, ensuring unique representations for each entity type via learning independent weight matrices for each type (Hu *et al*. 2020). This distinguishes HGT from other attention-based GNN architectures, such as graph attention networks (GAT) (Veličković *et al*. 2017). HGT aggregates and integrates information from source nodes to generate meaningful representations for target nodes through three key sub-modules within each HGT block (Figure 1e).

#### Attention Mechanism

The first sub-module computes the attention score between source nodes *(s)* and target nodes *(t)* connected by an edge. Inspired by the Transformer architecture (Vaswani *et al*. 2017), the source and target node vectors are mapped to Key and Query vectors, respectively, using type-specific linear projections. To account for heterogeneity, each node type has an independent linear projection, ensuring unique modelling of all node types. The similarity between the Key and Query vectors is calculated using an edge-type-specific attention matrix (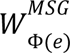), where Φ(*e*) denotes the edge type. This approach enables the model to capture semantic differences across edge types, even when multiple edge types connect the same pair of nodes. Additionally, a tensor *(μ)* is introduced to encode the importance of each node-edge-node triple type, weighting their contributions to the target node. The final attention vector for a node pair is obtained by concatenating the attention heads calculated for that pair.

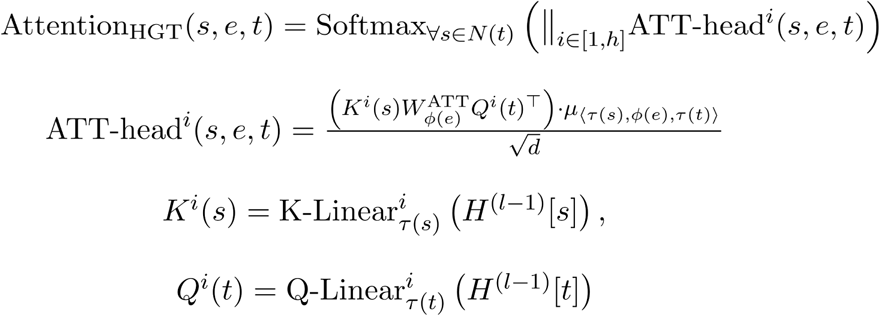

*s* represents the source node, *t* denotes the target node, and *e* is the edge connecting the source and target nodes. The set of all neighbouring nodes of *t* is represented as *N(t)* and *h* denotes the number of attention heads. For attention head *i*, *K^i^(s)* and *Q^i^(t)* represent the key and query vectors for the source and target nodes, respectively. The key linear projection for the source node type in attention head *i* is denoted by *K-Linear_τ(s)_^i^,* while *Q-Linear_τ(t)_^i^* represents the query linear projection for the target node type in attention head *i*. The attention matrix for the edge type *Φ(*e*)* is given by 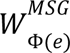 and *d* is the dimension of the linear projection vector. The importance tensor for the source node type, edge type, and target node type triple is denoted by *μ*. Finally, *H^(l-1)^[s]* and *H^(l-1)^[t]* represent the embedding vectors of the source and target nodes, respectively, from the previous layer.

#### Message Passing

In parallel with the attention score calculation, message passing occurs from source nodes to target nodes. The source node vector undergoes a linear projection to create a message vector, which is then transformed using an edge-type-specific message matrix 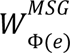. This ensures that the message-passing process captures the unique semantics of the edge type connecting the source and target nodes.

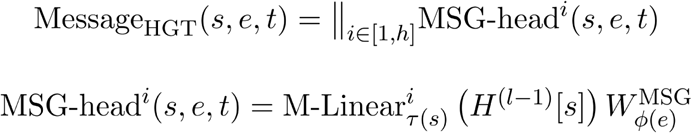

The message linear projection for the source node type in message head *i* is represented by *M-Linear_τ(s)_^i^*, while 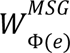 denotes the message matrix for the edge type *Φ*(*e*). Lastly, *H^(l-1)^[s]* is the embedding vector of the source node at the previous layer.

#### Feature Aggregation and Node Update

After attention scores and message vectors are computed, the target node aggregates messages from all source nodes using the attention scores as weights. This aggregation integrates feature information from all neighbouring nodes into a single, updated representation for the target node. The aggregated vector is then passed through a non-linear activation function, a residual connection, and a final linear projection specific to the target node type, resulting in the updated node representation (*H^(l)^[t]*) for the current HGT block (*l*).

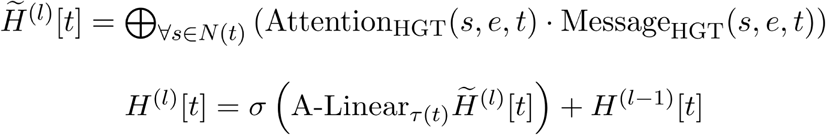

The set of all neighbouring nodes of *t* is represented as ***N(t)***, while ***σ*** indicates the non-linear activation function. The linear transformation specific to the type of the target node *t* is denoted by ***A-Linear_τ(t)_***. Lastly, ***H^(l-1)^[s]*** represents the embedding vector of the source node at the previous layer.

By stacking multiple HGT blocks, information propagates across the graph, enabling each node to capture not only its local context but also its relationships with distant nodes. ProtHGT models protein function prediction as a link prediction task, where the associations between protein nodes and Gene Ontology (GO) term nodes are inferred. For each potential protein-functional term node pair, the updated embeddings from the last HGT layer are concatenated and passed through a series of multi-layer perceptron (MLP) layers. These MLP layers transform the concatenated embeddings to capture complex interactions between the protein and functional term features. The output of the final MLP layer is passed through a sigmoid activation function to predict the probability of a relationship existing between the protein and the functional term.

### 2.3 Model Training and Implementation

The model was implemented using HGTConv convolution layers, the heterogeneous graph transformer (HGT) architecture provided by the PyTorch Geometric library (Fey and Lenssen 2019). Given the differences in scope, information content, and structure among the three Gene Ontology (GO) sub-ontologies (molecular function, biological process, and cellular component), an independent model was trained and evaluated for each sub-ontology.

For training, we employed a negative sampling strategy to approximate real-world conditions (i.e., a protein is annotated to only a few GO terms; therefore, most GO terms are non-associated with that protein). For each positive protein-GO term pair, we randomly selected 100 negative pairs from those not currently known to be associated. This strategy allowed the model to better differentiate between associated and non-associated pairs during training, improving its ability to learn meaningful relationships.

Model parameters were optimised using the binary cross-entropy loss function within an end-to-end learning framework. Since negative sampling introduced significant class imbalance, the loss function was weighted to amplify the contribution of positive predictions proportional to the negative sampling rate.

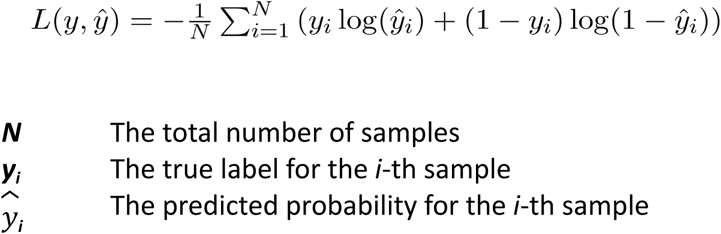

The dataset was split at the protein level into 80% training, 10% validation, and 10% test sets for training, hyperparameter tuning and performance evaluation, respectively. To ensure a realistic evaluation, we structured the graphs so that all edges associated with a given protein appeared exclusively in the training, validation, or test graph. This approach mimics real-world scenarios where newly sequenced proteins often lack relational information in databases. By preventing the model from seeing any relationship data for the test proteins during training, we aimed to simulate this cold-start situation and assess the model’s ability to predict functions for proteins with no prior connections.

A random search was conducted over the following hyperparameters: hidden channel size (16, 32, 64, 128, 256), learning rate (0.0001, 0.001, 0.005, 0.01), number of attention heads (2, 4, 8, 16), number of convolution layers (1, 2, 3, 4), batch size (512, 1024, 2048), neighbour sampling size (32, 128, 256), and weight decay (0, 1e-5, 5e-5, 0.0001, 0.001). The MLP layers used in the prediction module were also tuned, with hyperparameters including the dropout rate (0.2, 0.3, 0.4, 0.5), learning rate (0.0001, 0.001, 0.005, 0.01), and weight decay (0, 1e-5, 5e-5, 0.0001, 0.001). Through this tuning process, we identified the best-performing model configurations, specifically adapted to the characteristics of each GO sub-ontology.

### 2.4 Evaluation

The evaluation of ProtHGT and its variations, such as ablation models, was conducted on the 10% test dataset allocated during the initial data split. To further validate the model, its performance was compared with similar methods using temporal/time-based benchmark datasets. The first was the CAFA3 benchmark dataset (Zhou *et al*. 2019), the most up-to-date dataset with fully published results from the Critical Assessment of Functional Annotation (CAFA) competition. In CAFA3, the target protein list for function prediction was released in September 2016, and competing methods were trained on the latest available database versions and annotations up to that time. Predictions were generated prior to the annotation collection period, which spanned from February to November 2017, and experimentally validated annotations collected during this period served as the ground truth for evaluating prediction performance.

Recognising that the CAFA3 dataset is dated and does not include comparisons with recent graph-based models, ProtHGT was also evaluated on the time-based split test dataset introduced in the DeepHGAT study (Zhao *et al*. 2024). This dataset follows the same methodology as the CAFA benchmark but incorporates more recent annotations. Specifically, all functional annotations available for three species (Human, Mouse, and Arabidopsis) prior to 2021-09-01 were used for training, while annotations curated between 2021-09-01 and 2023-04-01 formed the test set. This setup allowed for a more up-to-date evaluation of ProtHGT and enabled comparisons with state-of-the-art graph-based methods. We also made sure that the rest of our source data obeys the time-based splits of the CAFA3 and DeepHGAT datasets.

The performance results of other methods on these datasets were sourced directly from their respective publications. To ensure a fair comparison, separate training and test graphs were constructed to align with the timelines of the benchmark datasets. Protein-GO term edges were first removed from the original graph described in Section 2.1, and then protein-GO term edges corresponding to the specific date ranges of each benchmark dataset were added to create time-specific graphs. These graphs were then subjected to independent model optimisation and evaluation. For the CAFA3 dataset, predictions were generated for every possible combination of proteins and GO terms included in the ground truth set. Protein-GO term pairs present in the ground truth were evaluated as positive, while those absent were treated as negative. For the DeepHGAT test set, pre-defined positive and negative labels for protein-GO term edges were used directly for evaluation.

#### Performance Metrics

To evaluate the models, we employed three metrics that are widely used in binary classification and link prediction tasks. The first is the F1-score, which is the harmonic mean of precision (pr) and recall (rc). The area under the precision-recall curve (AUPR) measures the trade-off between precision and recall, providing a comprehensive evaluation by quantifying the area under the precision-recall curve. Additionally, the Matthews Correlation Coefficient (MCC) evaluates the overall prediction performance while accounting for class imbalance. This makes MCC particularly valuable for cases like function prediction, where negative samples significantly outnumber positive ones.

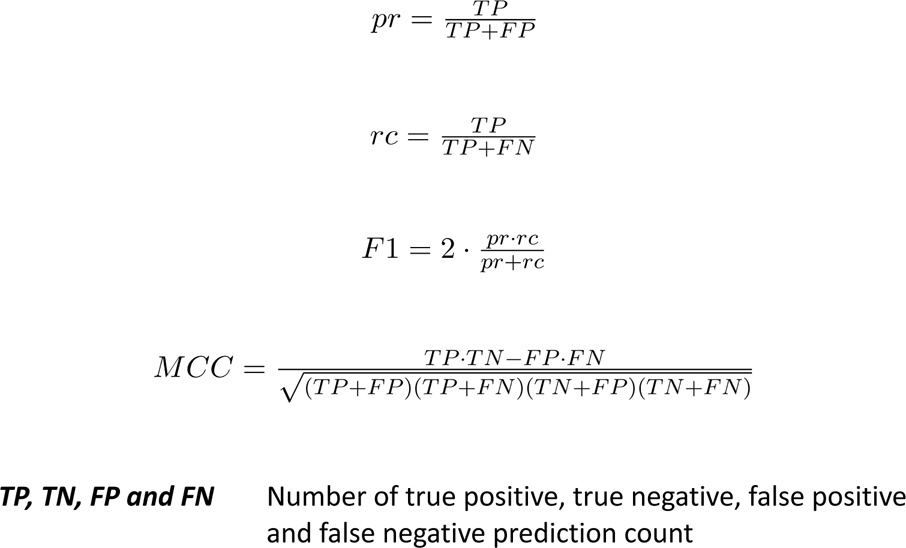

When comparing ProtHGT to other methods in the CAFA benchmark, we used their protein-centric evaluation mode metrics. This evaluation mode measures the accuracy of GO term predictions for each protein using two key metrics: Fmax and Smin. Both metrics rely on probabilistic predictions generated by the model across varying threshold levels, where predictions exceeding the threshold are classified as positive and those below it as negative.

The first metric, Fmax, represents the highest F1-score achieved across all threshold levels. The second metric, Smin, captures the minimum semantic distance between true annotations and predictions. It is calculated using two components: remaining uncertainty (ru) and misinformation (mi). These components incorporate the information content of GO terms, ensuring that predictions for less informative terms contribute less to the evaluation compared to predictions for more informative terms (Zhou *et al*. 2019).

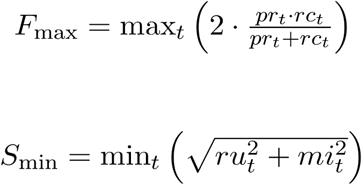

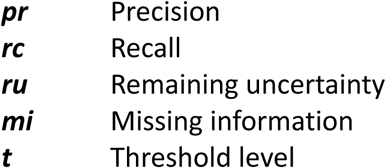

## 3 Results & Discussion

### 3.1 Performance Comparison with Other Methods

#### Evaluation on the DeepHGAT Test Set

To comprehensively evaluate the performance of ProtHGT, we compared it with a range of existing methods using the time-based split dataset introduced in the DeepHGAT study (Zhao *et al*. 2024). DeepHGAT is a graph attention network designed to integrate multiple biological data sources, such as protein-protein interaction (PPI) networks, ontology structures and protein sequences. By leveraging relationship-specific attention mechanisms, DeepHGAT captures both biological and structural context.

For consistency and comparability, the performance results of the other methods on this dataset were directly obtained from the original DeepHGAT study. The methods included in the comparison span a wide range of approaches and categories. PPI network-based models, including MV (Schwikowski, Uetz and Fields 2000), ProPN (Guangyuan *et al*. 2016), IFDR (Yu *et al*. 2017), DeepGraphGO (You *et al*. 2021), and PSPGO (Wu *et al*. 2023), utilise protein interaction data to infer annotations. MV serves as a baseline method, applying a straightforward “Guilt by Association” rule to predict annotations based on neighbour voting within the PPI network. IFDR enhances performance by employing random walks to uncover latent relationships and applying singular value decomposition (SVD) to reduce noise. DeepGraphGO and PSPGO, on the other hand, combine sequence feature extraction using the InterProScan model with graph neural network (GNN) approaches for PPI propagation. Among these, PSPGO stands out for its use of graph attention networks (GATs) integrated with label propagation.

Sequence-based models, such as PANDA2 (Zhao, Liu and Wang 2022), TALE+ (Cao and Shen 2021), PFresGO (Pan *et al*. 2023), and DeepFRI (Gligorijević *et al*. 2021), rely on pre-trained language models to extract sequence features. PANDA2, TALE+, and PFresGO integrate Gene Ontology (GO) structure relationships into their predictions. Among them, PFresGO stands out with its use of a graph attention framework, which captures and leverages the intricate relationships within the GO structure. DeepFRI takes a different approach by relying on protein structure data rather than PPI networks.

The results in Table 3 demonstrate that ProtHGT, particularly when using ProtT5 embeddings, outperforms all other methods across every evaluation metric and GO sub-ontology. Pre-trained on extensive protein sequence datasets, ProtT5 captures both local sequence patterns and global contextual relationships, enabling it to provide highly informative input features for downstream predictions. While TAPE and ESM2 embeddings also deliver strong results, they fall slightly behind ProtT5, likely due to differences in pre-training objectives, dataset sizes, or model architectures. Nevertheless, all three pre-trained protein language models offer significant advantages in protein function prediction, as they provide rich sequence-level representations that establish a robust foundation for further learning within the ProtHGT framework.

**Table 3.** Performance comparison with similar methods on the DeepHGAT Human dataset across GO sub-ontologies (the best performance for each metric and GO sub-ontology is shown with bold font)

|  |  | MV | ProPN | IFDR | DeepFRI | PANDA2 | TALE | PFreeGO | DeepGraphGO | PSPGO | DeepHGAT | ProthGHT-ESM2 | ProthGHT-TAPE | ProthGHT-ProtT5 |
| --- | --- | --- | --- | --- | --- | --- | --- | --- | --- | --- | --- | --- | --- | --- |
| Micro<br>AvgF1 | CC | 81.69 | 83.86 | 84.87 | 78.54 | 83.32 | 79.22 | 85.33 | 79.05 | 84.41 | 86.81 | 88.72 | 89.26 | **91.27** |
|  | MF | 80.82 | 74.27 | 81.52 | 79.19 | 78.06 | 81.97 | 81.97 | 81.30 | 82.39 | 83.60 | 85.60 | 86.06 | **89.33** |
|  | BP | 76.91 | 78.42 | 79.39 | 75.63 | 78.94 | 77.66 | 79.83 | 76.62 | 79.28 | 81.74 | 79.13 | 88.81 | **90.09** |
| Macro<br>AvgF1 | CC | 77.10 | 79.76 | 79.98 | 76.70 | 79.02 | 85.66 | 80.12 | 81.81 | 81.19 | 85.10 | 88.72 | 89.23 | **91.27** |
|  | MF | 74.17 | 79.96 | 80.55 | 78.35 | 81.14 | 82.80 | 82.34 | 80.99 | 82.48 | 85.89 | 85.53 | 86.02 | **89.32** |
|  | BP | 76.76 | 85.12 | 85.20 | 83.05 | 85.10 | 85.85 | 83.56 | 83.60 | 84.82 | 86.90 | 78.52 | 88.80 | **90.09** |
| AUC | CC | 76.70 | 81.98 | 82.66 | 81.90 | 83.54 | 86.57 | 88.55 | 85.02 | 86.93 | 89.29 | 95.35 | 96.76 | **97.16** |
|  | MF | 76.79 | 83.97 | 84.14 | 80.95 | 83.97 | 85.17 | 85.97 | 86.37 | 86.95 | 87.07 | 93.23 | 92.55 | **95.40** |
|  | BP | 76.15 | 87.05 | 88.05 | 81.45 | 82.73 | 83.31 | 85.78 | 84.08 | 86.83 | 88.39 | 88.96 | 95.54 | **96.58** |
| AUPR | CC | 56.47 | 84.76 | 85.71 | 81.76 | 83.13 | 84.51 | 88.20 | 85.13 | 89.85 | 90.32 | 95.25 | 96.74 | **97.14** |
|  | MF | 55.41 | 84.74 | 85.42 | 79.50 | 81.89 | 82.62 | 84.37 | 82.55 | 85.72 | 87.84 | 92.64 | 93.40 | **95.80** |
|  | BP | 58.82 | 83.52 | 82.67 | 76.92 | 78.82 | 79.19 | 81.56 | 80.25 | 82.28 | 85.17 | 90.06 | 95.27 | **96.62** |
| Fmax | CC | 73.32 | 78.88 | 79.06 | 77.16 | 78.21 | 79.05 | 81.50 | 79.37 | 81.93 | 83.83 | 88.58 | 88.64 | **91.12** |
|  | MF | 72.89 | 79.11 | 79.60 | 76.33 | 78.63 | 77.44 | 79.30 | 78.68 | 79.39 | 80.58 | 84.49 | 85.21 | **88.95** |
|  | BP | 69.06 | 79.15 | 78.79 | 76.81 | 78.07 | 78.00 | 78.60 | 77.58 | 78.16 | 79.28 | 74.89 | 88.37 | **90.05** |

Other methods in the comparison, such as DeepHGAT and PSPGO, demonstrate strong performance but are generally outperformed by ProtHGT. This superiority can be attributed to ProtHGT’s ability to leverage a highly heterogeneous graph structure that integrates a broader range of biological datasets, including pathway involvement and domain content. By incorporating these diverse sources, ProtHGT constructs a richer and more comprehensive representation space, enabling more accurate and biologically meaningful function predictions. In contrast, methods like DeepHGAT and PSPGO rely primarily on protein-protein and protein-functional term relationships, without fully exploiting proteins’ connections to other biological entities, potentially limiting their predictive capabilities.

Sequence-based models, such as PFresGO, PANDA2, and TALE+, fall behind ProtHGT despite utilizing advanced language models for protein sequence data. Their primary limitation again lies in the lack of integration with other biological relationships, which are crucial for capturing a broader and more nuanced biological context. Similarly, DeepFRI, which focuses on protein structure data, performs poorly compared to most methods. This underperformance is likely due to its reliance on coarse-grained sequence feature extraction, which lacks the granularity and depth potentially required for accurate protein function predictions.

#### Results on the CAFA3 Benchmark

To further evaluate ProtHGT, we compared its performance with other protein function prediction methods using the CAFA3 benchmark dataset (Zhou et al. 2019). While CAFA3 focuses on earlier machine learning and homology-based approaches and does not include comparable graph-based methods, we included it in our study due to its significance and widespread recognition in protein function prediction research. The evaluation was conducted in “full mode”, encompassing all no-knowledge proteins—test proteins that lacked annotations before the dataset’s release. To investigate the role of different protein representations, we tested both traditional embedding methods, such as APAAC, and modern pretrained protein language models, including ProtT5, TAPE, and ESM2. ProtHGT variants were compared against the top 10 models from the CAFA3 competition for each functional term category, along with two baseline methods: Naive and BLAST. The performance of all ProtHGT variants was evaluated using the Fmax and Smin metrics, as illustrated in Figure 2.

**Figure 2.**
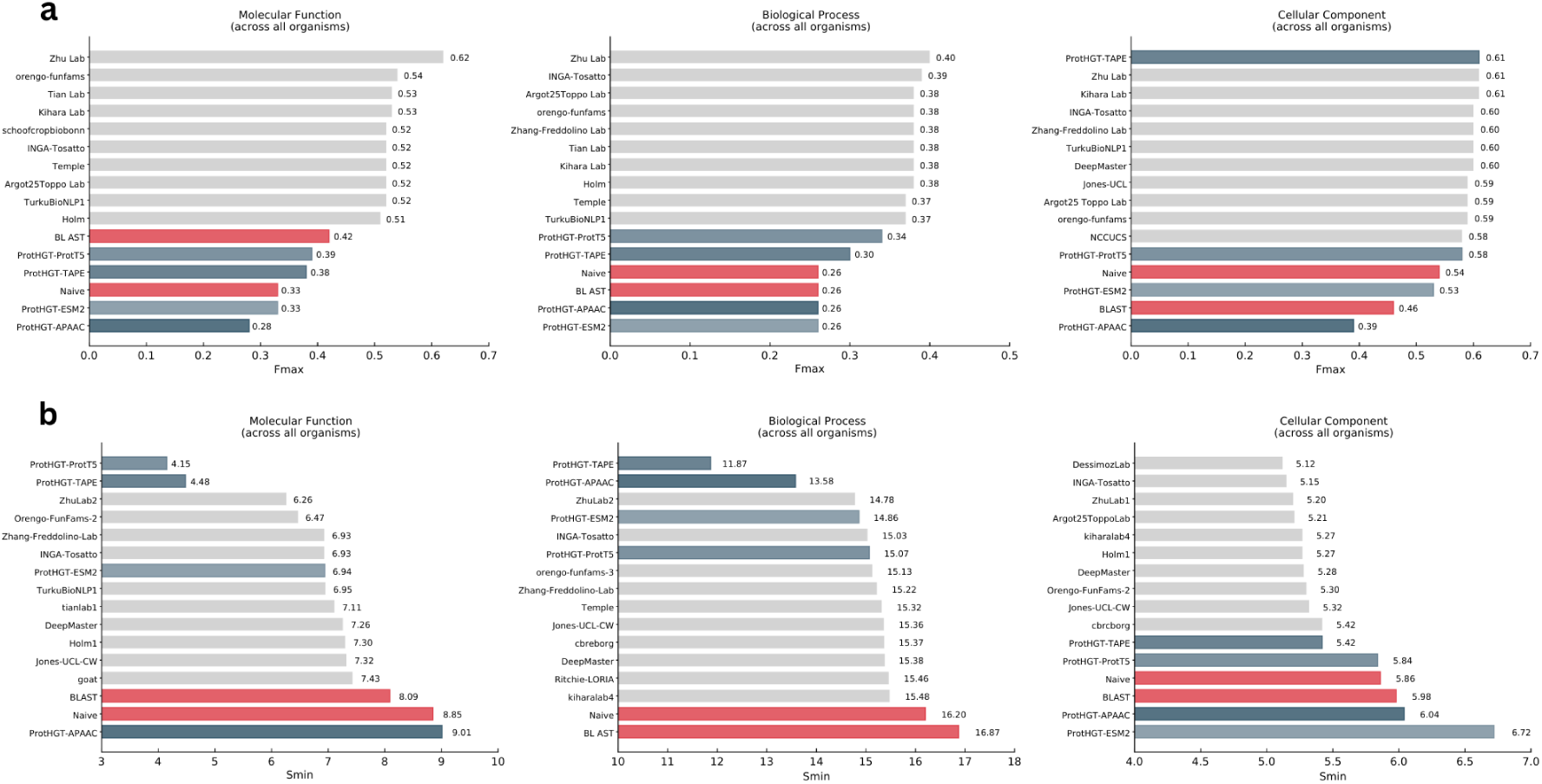
Performance comparison of ProtHGT models—ProtHGT-ProtT5, ProtHGT-TAPE, and ProtHGT-ESM2—against the top 10 performing methods (light grey) and two baseline methods (red) on the CAFA3 benchmark, evaluated using a) Fmax and b) Smin metrics. Higher Fmax scores indicate better accuracy in predicting protein functions, while lower Smin scores signify better performance in identifying highly specific and informative terms. For clarity, the performance scores for each method are displayed to the right of the corresponding bars.

ProtHGT consistently outperformed baseline methods, Naive and BLAST, across all GO sub-ontologies, underscoring its effectiveness as a graph-based protein function prediction model. Particularly in the Molecular Function (MF) and Biological Process (BP) categories, ProtHGT variants achieved top Smin scores (Figure 2b). The Smin metric evaluates a model’s ability to predict highly specific and detailed GO terms by incorporating their information content, assigning greater weight to precise predictions of informative terms over general ones. ProtHGT’s consistently low Smin values in the MF and BP sub-ontologies highlight its ability to prioritise and accurately predict these fine-grained terms. This suggests that ProtHGT not only captures general patterns but also provides biologically meaningful annotations that reflect the specific roles of proteins in these sub-ontologies.

In the Cellular Component (CC) category, ProtHGT achieved the highest Fmax score (Figure 2a), demonstrating its strength in accurately identifying broad protein associations within this sub-ontology. However, this was accompanied by slightly higher Smin values compared to baseline methods (Figure 2b), reflecting a trade-off in its ability to predict highly specific, information-rich terms. This trade-off suggests that while ProtHGT excels at capturing general protein associations in CC, further refinement is needed to improve its precision for highly specific cellular component terms. One likely explanation for this limitation is the sparse experimental annotations available in the dataset for features such as domain content, which play a critical role in identifying protein localisation. This lack of dense annotations constrains the model’s ability to uncover detailed functional relationships for high-information-content CC terms. Despite this, ProtHGT’s strong and consistent performance across all sub-ontologies highlights its robustness and adaptability. It is worth noting that the CAFA3 benchmark only publishes the results of the top 10 performing methods selected from a much larger pool of competing approaches. ProtHGT’s ability to achieve results that are on par with these top methods showcases its competitiveness.

When evaluating prediction models, it is important to consider not only their performance but also their usability and interpretability. Many high-performing methods, such as those featured in CAFA3, are often complex and computationally intensive, making them difficult to adapt for use outside the specific challenge context. Additionally, these sophisticated approaches frequently lack interpretability, making it hard for researchers to understand the reasoning behind specific predictions. The absence of user-friendly tools or reproducible programmatic frameworks further limits their practical application in biological research (Ulusoy and Doğan 2024). ProtHGT combines strong predictive accuracy with usability and interpretability, offering clear insights through its attention-based architecture. The use of heterogeneous graph data, which captures complex biological relationships between nodes, further enhances the interpretability of its predictions by revealing the underlying connections that drive its outputs. Additionally, its accessibility as both a programmatic tool and user-friendly web service ensures it is practical and adaptable for researchers across various applications beyond benchmarks.

The results also emphasise the critical role of embedding choice in determining prediction accuracy. Among the ProtHGT variants, ProtHGT-ProtT5 consistently achieved the best results across all metrics and sub-ontologies. Generally, pretrained protein language models demonstrated an ability to encode richer and more relevant protein features, significantly enhancing graph-based function prediction. In contrast, the traditional APAAC embedding lagged behind in performance.

Overall, these findings validate the effectiveness of ProtHGT’s integration of diverse biological data and advanced embeddings, establishing it as a robust tool for protein function prediction and pointing to areas for future refinement.

### 3.2 Ablation Analyses

#### Visualisation of Initial and Refined Embedding Spaces

To evaluate the biological validity of the ProtHGT representations (i.e., HGT-layer updated embeddings of proteins), we employed two widely used dimensionality reduction techniques, t-SNE (Van der Maaten and Hinton 2008) and UMAP (McInnes, Healy and Melville 2018), to transform the high-dimensional embeddings of proteins and GO terms into 2D visualisations (Figure 3). Specifically, this analysis aimed to assess whether the initial protein embeddings accurately represent biological attributes and whether the application of HGT enhances the learning of a protein’s inherent characteristics.

**Figure 3.**
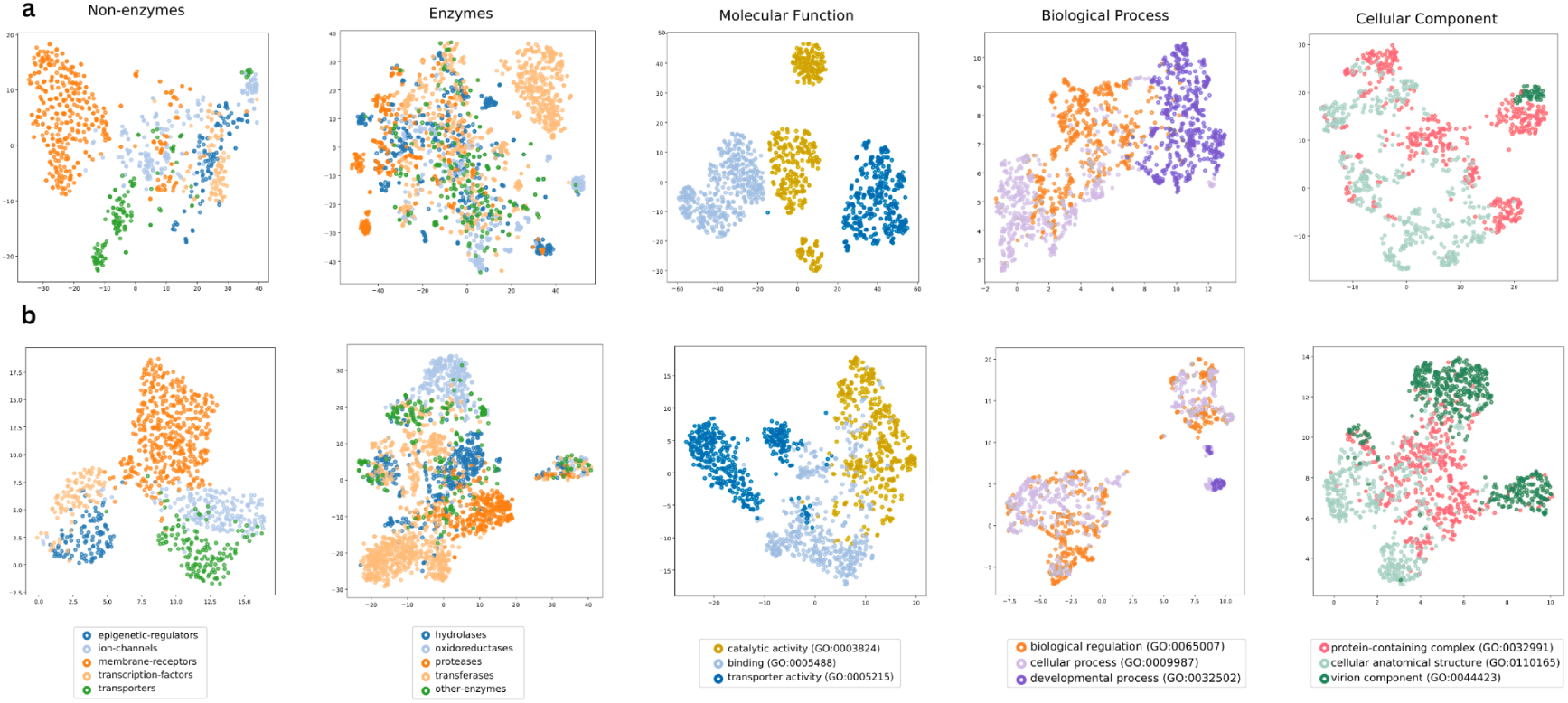
Protein and GO term embedding spaces before and after HGT refinement, visualised using dimensionality reduction techniques (UMAP and t-SNE) **a)** Initial embedding spaces for proteins (TAPE embeddings), grouped by protein superfamilies, and GO terms (anc2vec embeddings), organised by functional categories. **b)** Refined embedding spaces, i.e., ProtHGT embeddings, obtained via passing initial embeddings through HGT layers during the ProtHGT model training, showing improved clustering and separation of biologically meaningful groups.

The analysis was conducted using a dataset of proteins, which were categorised into five enzyme superfamilies (hydrolases, oxidoreductases, proteases, transferases, and other enzymes) and five non-enzyme superfamilies (epigenetic regulators, ion channels, membrane receptors, transcription factors, and transporters) (Atas Guvenilir and Doğan 2023). In t-SNE and UMAP projection plots (Figure 3), proteins are colour-coded by their superfamilies, allowing us to assess whether proteins from the same family formed cohesive clusters in the embedding space. The clustering of proteins within their respective superfamilies reflects the ability of the embeddings to encode meaningful biological features that differentiate protein types.

For GO term projections, the child terms of second-layer terms (those directly below the root terms) were grouped together to analyse the organisation of the GO embedding space. From the second-layer terms, the three terms with the highest number of child terms were selected within each GO category, and 500 random child terms were sampled for visualisation. This sampling strategy reduced overcrowding in the figures while preserving a representative set of terms to evaluate the embedding structure.

The visualisations reveal that both the initial and refined embeddings effectively capture biologically meaningful relationships. Remarkably, the initial embeddings already exhibited strong clustering, particularly for non-enzyme proteins and molecular function GO terms (Figure 3a). Selecting high-quality initial feature vectors plays a crucial role in accurately representing the inherent characteristics of nodes. High-quality initial embeddings establish a strong foundation for downstream learning by providing contextually rich and biologically relevant features.

The HGT-refined embeddings further improved clustering, with enhanced separation and organisation of entities (Figure 3b). This improvement is particularly evident for non-enzymes, where distinct clusters emerge for different families.

Additionally, the HGT-enhanced embeddings improved the differentiation of GO terms within the cellular component category, although the clustering for both biological process (BP) and cellular component (CC) categories still remained less distinct. The less visible clustering in BP and CC categories can be attributed to several factors. BP terms often represent interconnected and overlapping pathways, making clear separation challenging. Similarly, CC terms frequently share localisation, as proteins can reside in multiple compartments, leading to overlap in their embeddings.

Overall, the findings emphasise the model’s ability to integrate structural and biological context through HGT layers, refining the initial representations into embeddings that are more biologically coherent and meaningful.

#### Impact of Node Types on Predictive Performance

To evaluate the contribution of different node types and their associated edges to model performance, we conducted two ablation analyses across the three Gene Ontology (GO) sub-ontologies. To focus solely on the impact of edges and their associated information on predictions, these analyses were performed on models with a simple dot product operation for the prediction module. This approach avoided the added representational and feature interaction capabilities of multi-layer perceptron (MLP) layers, which can amplify certain features or relationships and potentially introduce biases to the ablation analysis.

In the first experiment, we trained models on datasets where a specific node type and all its associated edges were removed. The results, shown in Figure 4a, reveal the effect of these removals on MCC across the three GO sub-ontologies. For the molecular function category, the highest performance is achieved with an MCC score of 0.697 when all node types are included, emphasizing the benefits of integrating diverse node and edge types into the analysis. Among all node types, removing EC number nodes caused the most significant performance drop, reducing the MCC to 0.509. This sharp decline underscores the critical role of EC numbers in providing insights about proteins’ enzymatic activities and the biochemical reactions they participate in, making them highly informative for molecular function predictions.

**Figure 4.**
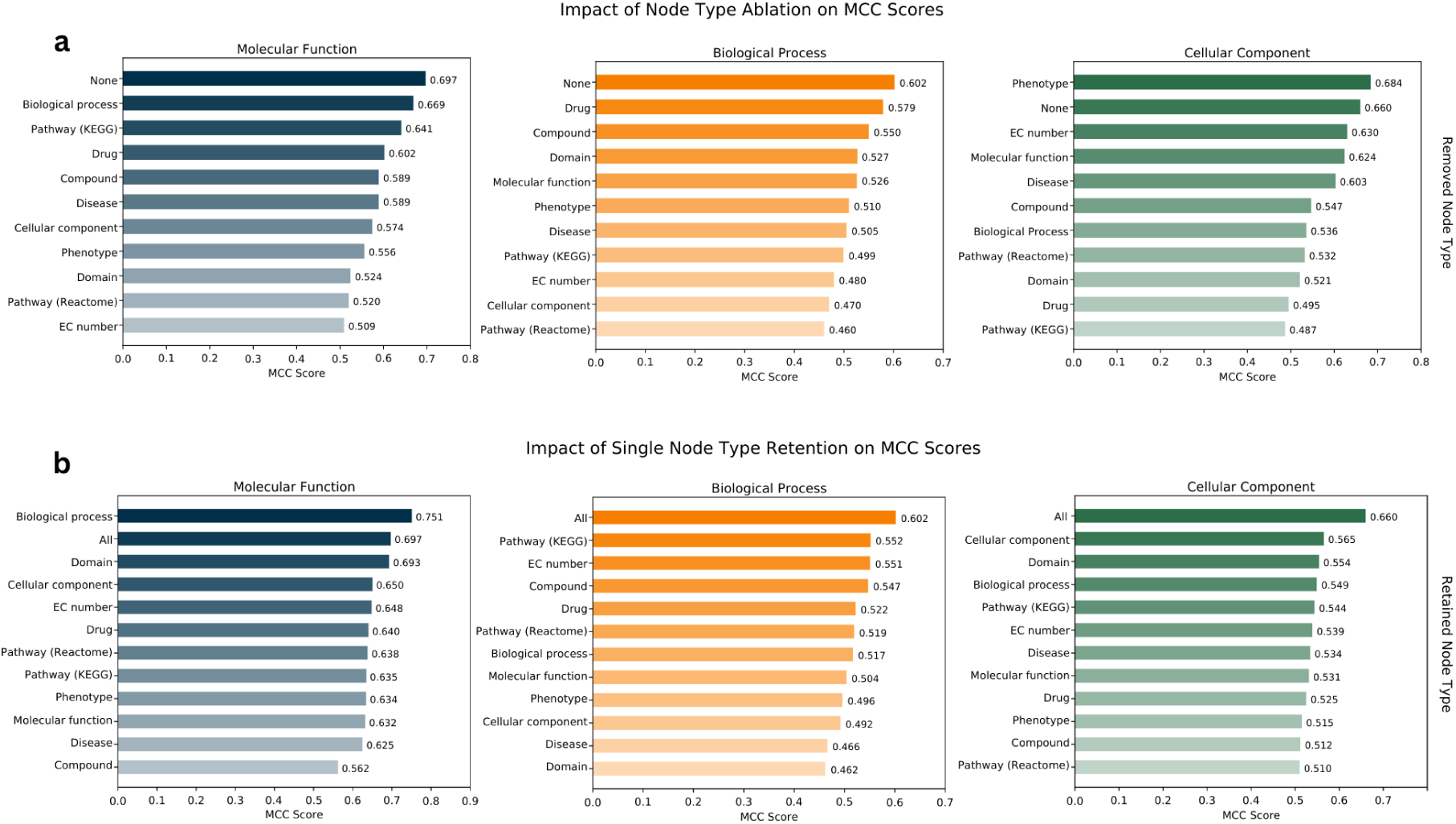
Impact of node type ablation and retention on MCC scores across Molecular Function, Biological Process, and Cellular Component sub-ontologies. **a)** MCC scores when individual node types are ablated, showing the effect of removing each node type from the graph. Lower MCC scores highlight the detrimental impact of excluding specific node types, with “None” representing the baseline where all node types are included. **b)** MCC scores when only a single node type is retained, illustrating the relative contribution of each node type to the model’s predictive accuracy. Higher MCC scores indicate a more substantial contribution of the retained node type, while “All” represents the baseline with all node types present.

In the biological process category, the highest MCC score of 0.602 was observed again with all node types retained. Similar to molecular function, the removal of EC number nodes caused a substantial performance drop, reducing the MCC to 0.480. These results further confirm the importance of EC numbers in capturing essential information required for functional annotation in biological process predictions. For the cellular component category, the highest MCC score of 0.684 was achieved when Phenotype nodes were ablated. This suggests that phenotype nodes may introduce excess complexity to the dataset when predicting cellular component terms, and their removal can simplify the graph structure in a way that benefits performance.

Notably, across all sub-ontologies, removing pathway nodes (whether from KEGG or Reactome databases) consistently caused significant performance declines. This highlights their critical role in providing contextual information about protein interactions within broader biological systems, such as metabolic or signalling pathways. Pathway nodes serve as connectors that enrich the overall graph structure by integrating proteins’ roles within complex biological networks. Conversely, the ablation of certain node types, such as Disease and Compound nodes, resulted in relatively smaller performance declines, suggesting that these nodes play a less prominent role in the specific functional contexts.

To examine the individual contributions of specific node types to training, we conducted a second ablation analysis. In this approach, models were trained on datasets where only a single node type and its associated edges were retained, while all other node and edge types were removed. This allowed us to isolate the impact of individual node types on model performance (Figure 4b).

For the biological process and cellular component categories, the highest performance was consistently achieved when all node types were retained, reaffirming the value of a holistic, multi-relational graph for protein function prediction. Interestingly, in the molecular function category, the model trained on a graph containing only biological process nodes achieved results close to those obtained with a graph containing all node types. This indicates that biological process annotations of proteins, as well as the hierarchical relationships within biological process terms and biological process-molecular function term associations, capture most of the critical information required for molecular function predictions. This finding implies that other node types, while valuable, play a secondary role in this specific sub-ontology.

Domain nodes also play a critical role in protein function prediction, particularly in the molecular function and cellular component sub-ontologies. Their removal leads to significant performance drops while retaining only domain nodes still achieves relatively strong predictions. As structural and functional units, domains directly influence a protein’s biochemical activities, such as enzymatic activity or binding specificity, and are often linked to cellular localisation through targeting signals or structural roles.

A particularly notable observation relates to pathway nodes. In the node removal ablation, their exclusion led to significant performance drops across all sub-ontologies, emphasising their importance in providing contextual information about protein interactions. However, in the node retention analysis, graphs containing only pathway nodes and their edges to proteins yielded moderate performance for molecular function and cellular component predictions. This indicates that pathway nodes derive much of their predictive power from their interactions with other node types, and their role appears to be integrative, serving as connectors that enhance the overall graph’s richness when combined with complementary nodes. In contrast, for the BP category, retaining only KEGG pathway nodes resulted in a relatively high MCC score of 0.552, suggesting that pathways can independently provide substantial functional insights in this sub-ontology, which was expected.

These results highlight the critical role of integrating diverse biological datasets for effective protein function prediction. While individual node types, such as biological process nodes for molecular function and pathway nodes for biological function prediction, can independently provide valuable insights, the collective use of multiple node types significantly enhances predictive performance across all GO sub-ontologies.

### 3.3 Evaluating the Biological Relevance of ProtHGT’s Predictions

To biologically validate ProtHGT’s predictions, we analyzed its predictions for randomly selected proteins. One particularly interesting case involved “Keratin-associated protein 23-1” (Uniprot ID: A1A580). ProtHGT correctly predicted both of this protein’s manual annotations from two distinct ontologies: “cytosol” (GO:0005829) from the cellular component ontology and “protein binding” (GO:0005515) from the molecular function ontology, with high probabilities of 0.996 and 0.999, respectively. These accurate predictions demonstrate ProtHGT’s ability to capture biologically relevant functions for this protein.

Additionally, ProtHGT predicted the function “structural constituent of bone” (GO:0008147) with a high probability (0.980), which was evaluated as a false positive since it does not appear in manually curated annotations or known databases. However, further investigation of ProtHGT’s training graph (Figure 5) revealed plausible reasons behind this prediction. The graph showed dense connectivity between Keratin-associated protein 23-1 and entities linked to bone-related functions. For example, the protein is associated with pathways such as “Keratinization” (R-HSA-6805567) and “Developmental Biology””

**Figure 5.**
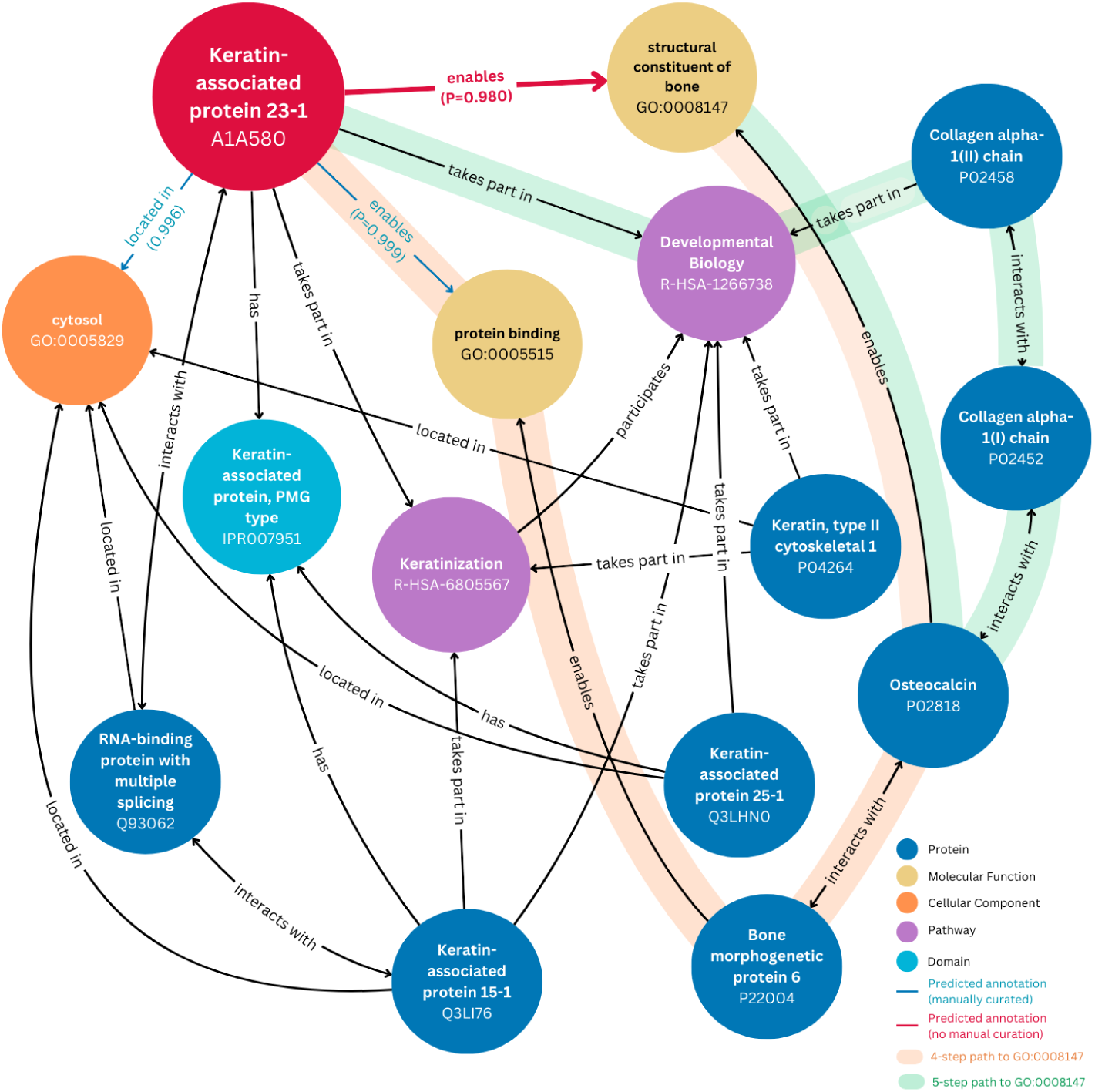
Key connections in the heterogeneous graph for the use case study of Keratin-associated protein 23-1 (A1A580) predictions. Probabilities for each predicted edge are displayed below the edge labels. Two paths linking A1A580 to the GO term “structural constituent of bone” (GO:0008147) are highlighted: a 4-step path in orange and a 5-step path in green.

(R-HSA-1266738), both of which are indirectly related to bone formation processes. Moreover, walks through the graph revealed connections via direct and indirect interactions with other proteins, such as “Collagen alpha-1(II) chain” (P02458) and “Bone morphogenetic protein 6” (P22004), which share common manually curated GO annotations and participate in overlapping pathways. The proteins involved in these pathways also share common domains, such as “Keratin-associated protein, PMG type” (IPR007951) further contributing to the indirect associations that likely influenced this prediction.

These findings indicate that ProtHGT can reveal functional overlaps or indirect associations that may not align with existing annotations but could point to previously uncharacterized biological roles for proteins. This opens opportunities for generating new hypotheses, such as potential links to disease mechanisms. This example highlights ProtHGT’s ability to combine predictive accuracy with interpretability and its potential to support hypothesis-driven research. By allowing researchers to trace the reasoning behind its predictions through its heterogeneous graph, ProtHGT facilitates the validation of known functions, exploration of unexpected functional links, and discovery of novel relationships, making it an invaluable tool for both prediction and discovery in functional genomics research.

## 4 Conclusion

In this study, we introduced ProtHGT, a heterogeneous graph transformer-based model for protein function prediction. By integrating diverse biological datasets—such as proteins, pathways, GO terms, EC numbers, domains, phenotypes, diseases, drugs/compounds, along with protein-protein interaction networks—ProtHGT constructs a comprehensive knowledge graph to capture complex biological relationships, enabling accurate predictions across all Gene Ontology (GO) sub-ontologies.

ProtHGT achieves state-of-the-art performance on multiple benchmarks, especially on the novel DeepHGAT dataset, consistently outperforming other methods by leveraging rich node feature vectors and integrating diverse biological contexts. Ablation analyses highlight the critical role of input protein language model embeddings along with heterogeneous data integration, showing that multiple node types, such as pathways and domains, significantly improve predictive accuracy. Embedding space visualisations further demonstrate ProtHGT’s ability to refine initial features into biologically meaningful representations. Finally, the use case study showcased ProtHGT’s ability to accurately capture known functions while also identifying potential novel associations through its graph-based reasoning. Additionally, it highlighted ProtHGT’s interpretability, allowing researchers to trace predictions back to their underlying connections, supporting both functional validation and hypothesis generation.

To ensure accessibility and usability, ProtHGT is provided as a programmatic tool hosted on https://github.com/HUBioDataLab/ProtHGT, complete with its models, source code, datasets and results. Additionally, a user-friendly web service is available on https://huggingface.co/spaces/HUBioDataLab/ProtHGT, allowing researchers with varying technical expertise to use ProtHGT easily.

Despite its strong performance, ProtHGT has some limitations that could be addressed in future work. One area for improvement is the model’s performance in sparsely annotated sub-ontologies, such as specific categories within the Biological Process (BP) or Cellular Component (CC) sub-ontologies. Exploring methods to better handle sparse annotations, such as transfer learning or leveraging self-supervised pretraining on unannotated protein data, could enhance predictive accuracy for these cases.

Additionally, while ProtHGT incorporates a broad range of node types, future work could investigate incorporating temporal data, such as context-dependent protein interactions or evolutionary trajectories, to capture dynamic properties of biological processes. Expanding the graph to include multi-omics data, such as transcriptomics or (epi)genomics, along with cellular information may further improve its applicability across diverse biological contexts. Lastly, future work could explore broader applications of ProtHGT, such as predicting gene/protein-disease associations, or modeling drug-target interactions for supporting drug discovery pipelines. These extensions would further validate the utility of ProtHGT in addressing a wide array of biological and biomedical challenges.

